# The IFT-A complex plays a major role in the assembly of anterograde intraflagellar transport trains

**DOI:** 10.64898/2026.07.03.736119

**Authors:** Adeline Mallet, Thierry Blisnick, Eloïse Bertiaux, Cécile Fort, Manuel Majrouh, Sylvain Trépout, Philippe Bastin

**Author notes:** equal contribution (Adeline Mallet, Thierry Blisnick). Ramaciotti Centre for Cryo-Electron Microscopy, Monash University, Monash, Victoria, Australia (Sylvain Trépout).

## Abstract

Cilia are assembled by intraflagellar transport (IFT), which relies on two protein complexes: IFT-A and IFT-B. It is generally assumed that IFT-B and IFT-A are critical for anterograde and retrograde transport, respectively. However, full deletion of *IFT-A* genes in several organisms suggests a possible contribution to anterograde transport. In many species, cilia collapse when IFT is altered, hindering functional studies. Here, we investigated the role of IFT-A in the protist *Trypanosoma brucei*, where IFT is not required for cilium maintenance. Following the inducible knockdown of IFT88 (an IFT-B member) or IFT140 (an IFT-A member), we monitored the fate of several IFT proteins in preassembled cilia using live imaging and evaluated the consequences on train formation by volumetric electron microscopy. Surprisingly, both IFT88 and IFT140 turned out to be essential for anterograde train assembly. Their depletion initially led to the formation of shorter trains and subsequently to an inhibition of train injection. We propose a model to reconcile the diverging phenotypes reported in the literature.

## Introduction

Cilia and flagella are a landmark feature of eukaryotes where they perform multiple functions such as motility, signalling or morphogenesis. They are constructed by the action of intraflagellar transport (IFT), the bidirectional movement of large protein particles (also called trains) taking place along axonemal microtubules underneath the membrane (Kozminski et al., 1993)(review in (Lacey and Pigino, 2025)). Motors of the kinesin 2 or the cytoplasmic dynein 2 families are responsible for anterograde and retrograde transport, respectively. The IFT dynein is transported in an inactive form as a cargo of anterograde trains before powering retrograde transport (Jordan et al., 2018). IFT proteins have been purified from flagella of the green alga *Chlamydomonas* and can be separated in two complexes termed the IFT-A (6 proteins) and the IFT-B (16 proteins) complexes (Cole et al., 1998). IFT proteins and motors are conserved in most species with cilia and flagella (van Dam et al., 2013). Both complexes have been reconstructed *in vitro* revealing their peculiar architecture (Hesketh et al., 2022; Jiang et al., 2023; Ma et al., 2023; Meleppattu et al., 2022; Petriman et al., 2022). Structural analysis by cryo-electron microscopy of the green alga *Chlamydomonas* revealed the organisation of anterograde IFT trains within the flagellum (Jordan et al., 2018). These trains measure on average 312 nm and contain several repetitive densities of 6, 11 and 18 nm. Using mutants for the IFT dynein heavy chain and the IFTA protein IFT139, the three densities could be assigned to IFT-B, IFT-A and dynein, respectively. Their ratio was estimated around 8 copies of IFT-B for 4 copies of IFT-A and 2 copies of dynein (Jordan et al., 2018). Furthermore, a combination of cryo-electron microscopy and of alphafold2 predictions unveiled the intimate structure of the IFT-A and IFT-B complexes and their interactions within the trains themselves, bringing insights about how the IFT-A complex can form multimers with the help of IFT-B (Lacey et al., 2023).

Inhibition of IFT-B blocks cilium formation whereas depletion of IFT-A results in the construction of tiny organelles filled with IFT material. This has been observed using mutant cell lines or RNAi knockdown experiments in multiple organisms ranging from animals (mammals, *Caenorhabditis elegans*, *Drosophila melanogaster*) to algae (*Chlamydomonas*) and protists (*Trypanosoma*, *Tetrahymena*)(Absalon et al., 2008b; Blacque et al., 2006; Han et al., 2003; Huangfu et al., 2003; Iomini et al., 2009; Tsao and Gorovsky, 2008). This led to a general consensus that IFT-B is essential for anterograde transport while IFT-A would be required for retrograde transport. However, several pieces of evidence indicate that IFT-A proteins could have more functions. First, deletion of the *IFT-A* genes *IFT121*, *IFT122*, *IFT144* (Zhu et al., 2017) or *IFT140* (Picariello et al., 2019) in *Chlamydomonas* resulted in absence of flagella. Second, knockout of the *IFT140* gene in the protist *Leishmania mexicana* (Sunter et al., 2018) also resulted in absence of flagellum. In all cases, little or no accumulation of IFT-B proteins were noticed, which does not match with a restricted function of IFT-A in retrograde transport. Therefore, different approaches can result in conflicting phenotypes, raising the need for further experimentation.

In most investigated species, cilia rely on IFT for their construction but also for their maintenance (Eguether et al., 2018; Kozminski et al., 1995), meaning that any interference with IFT results in collapse of the organelle, hindering detailed analysis. We circumvented this issue by monitoring IFT protein knockdown in the protist *Trypanosoma brucei*. This organism assembles a new flagellum at each cell cycle while maintaining the existing flagellum (Sherwin and Gull, 1989), hence allowing the examination of the organelle in its growing and maintenance state in the same cell.

*T. brucei* possesses the full complement of IFT proteins (Absalon et al., 2008b; Julkowska and Bastin, 2009; van Dam et al., 2013) and is amenable to genetic manipulation. Tagging of IFT proteins and motors displayed typical anterograde and retrograde intraflagellar transport (Bertiaux et al., 2018a; Blisnick et al., 2014; Buisson et al., 2013) and inhibition of IFT blocks flagellum construction (Davidge et al., 2006; Franklin and Ullu, 2010; Kohl et al., 2003). Initial studies revealed that knockdown of IFT122 or IFT140, two core components of the IFT-A complex, resulted in the construction of short flagella filled with electron dense elements resembling IFT trains, a finding confirmed by immunofluorescence staining of IFT-B proteins (Absalon et al., 2008a; Absalon et al., 2008b; Fort et al., 2016). Importantly, depletion of IFT protein by RNA interference (RNAi) knockdown inhibits only flagellum construction and not maintenance of flagellum length (Fort et al., 2016). This offers the opportunity to monitor the impact of IFT depletion over time in existing flagella that were assembled before the onset of RNAi.

Here, we used RNAi knockdown targeting IFT88 and IFT140 as representative of the IFT-B and IFT-A complexes, respectively. The impact of their depletion on IFT was monitored over time in flagella that had been assembled prior to RNAi. Two-colour live imaging was used to track simultaneously an IFT-A and an IFT-B protein and train length was measured by focused ion beam scanning electron microscopy (FIB-SEM). This analysis revealed that IFT88 is essential for the construction of IFT trains as expected but surprisingly, that IFT140 is also required. We propose that the IFT-A complex contributes to efficient IFT-B polymerisation and to train elongation and that its absence inhibits anterograde train assembly, while reduced amounts is sufficient to sustain the assembly trains too short to support full dynein transport, hence explaining the retrograde transport defect reported in multiple publications.

## Results

### The 6 *IFT-A* genes are conserved in *Trypanosoma brucei*

The 6 *IFT-A* genes (IFT43, IFT121, IFT122, IFT140, IFT144) are conserved in *T. brucei* (Julkowska and Bastin, 2009) and 5 of them contain the typical protein-protein interacting domains WD40 or TPR (Fig. 1)(Taschner and Lorentzen, 2016). However, they have been much less studied than members of the IFT-B complex. The available data indicate a role in retrograde transport for IFT122, IFT140 and IFT144 following their silencing by RNAi, resulting in a mixture of cells, either with a very short flagellum containing a high amount of IFT172, or with no visible flagellum (Absalon et al., 2008a; Absalon et al., 2008b). Preliminary localisation of the 6 IFT-A proteins was reported in TrypTag (http://tryptag.org/), a global project aimed at determining where each expressed protein of the *T. brucei* genome localizes in the trypanosome cell (Billington et al., 2023). They appear concentrated at the base of the flagellum and along the length of the organelle, a typical localization for IFT proteins (Absalon et al., 2008b; Buisson et al., 2013).

**Figure 1.**
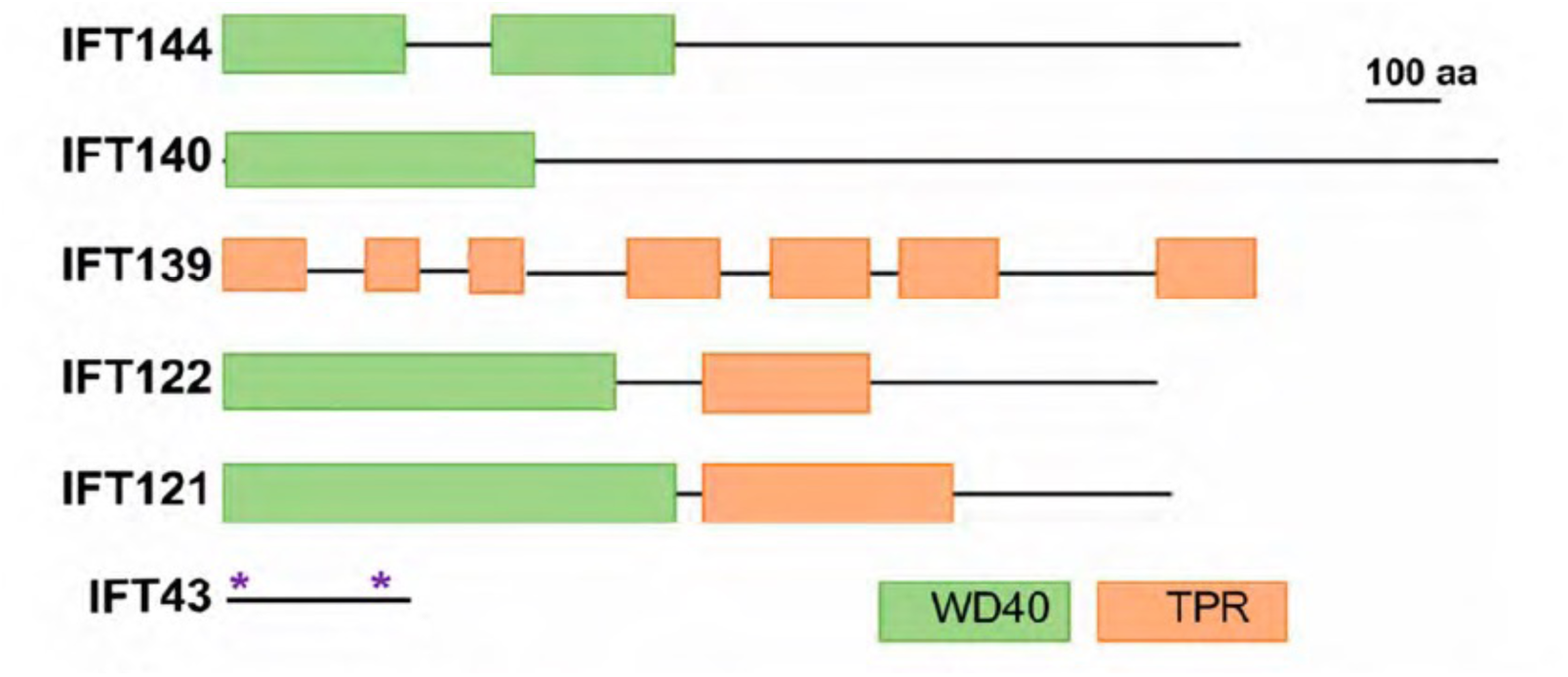
The 6 genes of the IFT-A complex are conserved in *T. brucei*. The major protein domains are shown in green (WD40) and orange (TPR, tetratricopeptide domain) for IFT144, IFT140, IFT139, IFT122 and IFT121. The two stars indicate domains of IFT43 that are not structured and could adopt different configurations.

### Localization and intraflagellar transport of IFT140, IFT 122 and IFT139

IFT is observed by live imaging of trypanosomes expressing a fluorescent reporter fused to the IFT protein of interest following endogenous tagging (Bertiaux et al., 2018a). We chose IFT122 and IFT140 as IFT-A core proteins and IFT139 for peripheral proteins. N-terminal labelling is preferred, as it enables control of expression by the 3’ untranslated region of the target gene, which is the main regulator of transcript abundance and turnover in trypanosomes (Clayton, 2002; Dean et al., 2015). Therefore, IFT140 and IFT139 were N-terminally tagged; however, IFT122 was tagged at the C-terminal as TrypTag reported better efficiency at this end (Billington et al., 2023). To enable functional studies, this was performed in a strain where IFT140 expression can be targeted by RNAi following tetracycline-inducible expression of double stranded RNA (Wang et al., 2000)(see below).

Tagging with mNeonGreen (mNG) revealed typical association to the base of the trypanosome flagellum and IFT movement for all three proteins (Fig. 2A-C and Video S1). Signal intensity appeared weaker compared to what was observed for IFT-B proteins (Bertiaux et al., 2018a; Bhogaraju et al., 2013), probably due to a lower concentration of complex A proteins in the flagellum. Indeed, proteomic analysis revealed that IFT-A proteins are about twice less abundant than IFT-B proteins in purified intact flagella (Subota et al., 2014). To quantify IFT trafficking, at least 20 videos of 30s were acquired for each cell line and kymographs were extracted (Fig. 2A-C, right panels). Over 200 anterograde trains were analysed and the frequency of injection (Fig. 2D) as well as the motion (Fig. 2E) of IFT trains were quantified. Anterograde trains labelled with mNG::IFT140, IFT122::mNG, and mNG::IFT139 exhibited similar average speed of 2-3 µm/s and frequency of 0.5 train/s under control conditions (Fig. 2D-E) (Table 1, non-induced samples). These values are comparable to data previously observed for other IFT-B proteins in an equivalent genetic context (Bertiaux et al., 2018a; Bhogaraju et al., 2013). Retrograde transport was regularly observed but the signals were too weak to be reliably quantified.

**Figure 2.**
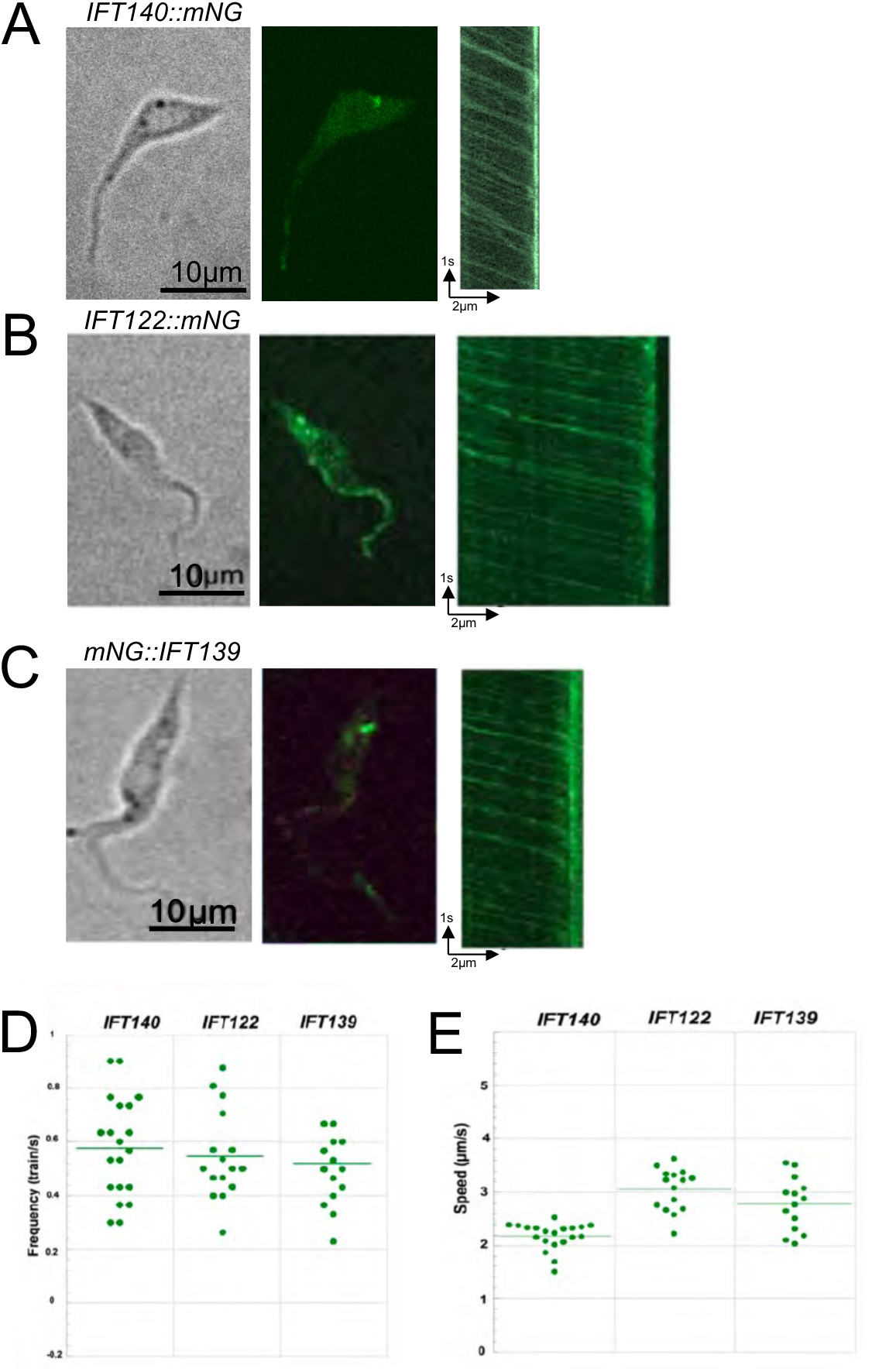
IFT-A proteins traffic in the trypanosome flagellum. A representative cell is shown expressing either (A) IFT140::mNG, (B) IFT122::mNG and (C) mNG::IFT139. (A-C) The left and central panels correspond to phase contrast and a still image of direct mNG fluorescence, respectively. Corresponding sequences are shown at Video S1. On the right panel, the kymographs represent the movement of (A) IFT140::mNG, (B) IFT122::mNG and (C) mNG::IFT139 over time. (D-E) Graphs representing anterograde IFT train frequency (D) and speed (E) measured in 20 different cells. Quantitative values are presented at Table 1 (N.I., non-induced samples, i.e. without RNAi knockdown).

**Table 1.** Frequency and speed of anterograde IFT trafficking of various IFT proteins in the *IFT140^RNAi^* cell line. A total of 20 films were analysed per condition. Frequency information and train speeds were extracted from 30s videos using Icy software and the Kymograph Tracker 2 plugin.

| ***IFT140*<sup>RNAi</sup> *mNG::IFT140*** | **Frequency (trains/s)** | **Speed (μm/s)** | **n trains** |
| --- | --- | --- | --- |
| NI | 0.53 ± 0.20 | 2.20 ± 0.45 | 353 |
| 24h | 0.36 ± 0.12 | 1.90 ± 0.53 | 296 |
| 36h | 0.31 ± 0.14 | 1.58 ± 0.58 | 141 |
| 48h | 0.05 ± 0.15 | 1.78 ± 0.46 | 17 |
| ***IFT140*<sup>RNAi</sup> *IFT122::mNG*** | **Frequency (trains/s)** | **Speed (μm/s)** | **n trains** |
| NI | 0.54 ± 0.16 | 3.10 ± 0.81 | 262 |
| 24h | 0.13 ± 0.12 | 1.30 ± 0.61 | 65 |
| 36h | 0.02 ± 0.05 | 1.24 ± 0.75 | 11 |
| 48h | 0.01 ± 0.06 | 1.65 ± 0.31 | 7 |
| ***IFT140*<sup>RNAi</sup> *mNG::IFT139*** | **Frequency (trains/s)** | **Speed (μm/s)** | **n trains** |
| NI | 0.52 ± 0.16 | 2.82 ± 0.81 | 235 |
| 24h | 0.25 ± 0.15 | 2.06 ± 0.79 | 128 |
| 36h | 0.09 ± 0.15 | 1.74 ± 0.57 | 63 |
| 48h | 0.03 ± 0.09 | 2.03 ± 0.55 | 22 |

### IFT122 and IFT139 depend on IFT140 for flagellar association and IFT transport

Following IFT140 silencing in the *IFT140^RNAi^* strain, the behaviour of the three fusion proteins was examined (Fig. 3). First, the fluorescence level of the mNG::IFT140 fusion protein was monitored as a reporter for RNAi efficiency (Fig. 3A-B). At intermediate induction times, the amount of IFT140 in the flagellum decreases and the frequency of labelled anterograde trains drops (Fig. 3A-B). Train speed in the uninduced condition is 2.2µm/s, and after induction, it is reduced between 1.6 and 1.9 µm/s (Table 1). After 48 h of knockdown, there is virtually no movement observed by live imaging in most cells and the flagella look much darker, proving the effectiveness of RNAi silencing (Fig. 3A, last kymograph).

**Figure 3.**
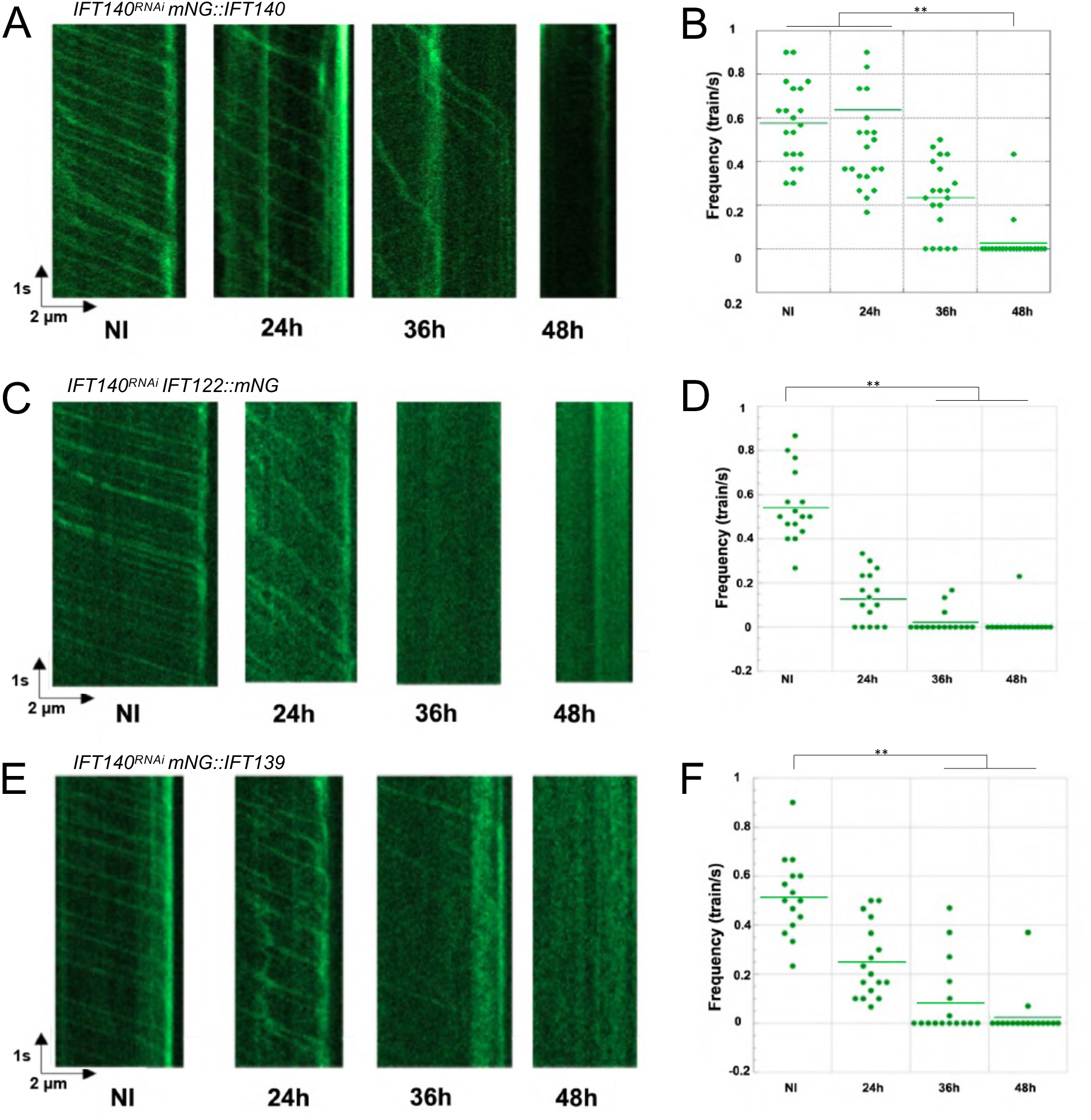
IFT140 is essential for trafficking of other IFT-A proteins. Representative kymographs extracted from videos acquired by fluorescent live cell imaging showing the evolution of IFT trafficking of the indicated proteins during the course of IFT140 knockdown (NI, RNAi non-induced) and graphical representations of the frequency of anterograde trains on (A,B) *IFT140^RNAi^ mNG::IFT140* (C,D) *IFT140^RNAi^ IFT122::mNG* and (E, F) *IFT140^RNAi^ mNG::IFT139* cell lines. The depletion of IFT140 results in arrest of trafficking of all three proteins. Quantitative values are shown at Table 1. ** p<0.05 (Anova test)

Next, we examined the effect of reduced IFT140 abundance on IFT122::mNG (Fig. 3C,D) and mNG::IFT139 (Fig. 3E,F) distribution. After RNAi induction, the anterograde transport frequency of IFT122::mNG and mNG::IFT139 decreases drastically. At intermediate time points in the RNAi kinetics, the speed of IFT122::mNG-tagged trains is reduced (Table 1). After 48 h of induction, there is virtually no movement of fluorescent signals detected in most flagella. Taken together, these results show that the transport of IFT122 and IFT139 is dependent on the presence of IFT140, as in other species (Behal et al., 2012; Mukhopadhyay et al., 2010) (Picariello et al., 2019) and validate its use as a reference for the IFT-A complex.

### IFT-A and IFT-B are always associated during anterograde transport

IFT trains have been described as multimers of IFT-A and IFT-B complexes transporting various cargoes. We investigated whether the trypanosome IFT-A proteins always travel with IFT-B proteins, or whether some trains contain only one or the other complex. We developed cell lines for observing IFT-A and IFT-B simultaneously under both control conditions and upon depletion of a protein from each complex. IFT140 with mNG tagging (Fig. 3A) and IFT81 with tandemTomato (tdT) fusion (Bertiaux et al., 2018b) were selected as representative reporters of the IFT-A and IFT-B complexes, respectively. For functional studies, the well-characterised cell lines *IFT88^RNA^*^i^ (Kohl et al., 2003) and *IFT140^RNAi^* (Absalon et al., 2008a) were transformed to express both mNG::IFT140 and tdT::IFT81. Cell lines were observed using a spinning disk microscope equipped with two cameras for simultaneous acquisition (Bertiaux et al., 2020). On the videos obtained, the fluorescent signals corresponding to mNG::IFT140 and tdT::IFT81 were easily visible (Video S2 and Fig. 4A for non-induced *IFT88^RNAi^* and Video S3 and Fig. 5A for non-induced *IFT140^RNAi^*). Fluorescent protein movements were visible in the flagellum for both green and pink channels, with a clear overlap between mNG::IFT140 and tdT::IFT81 (Fig. 4A and 5A, overlay). Kymograph extraction showed an almost perfect colocalization (Fig. 4A and 5A, left panels) and quantification revealed that 98% of anterograde trains contain both mNG::IFT140 and tdT::IFT81, supporting the view that all IFT trains are made of IFT-A and IFT-B complexes.

**Figure 4.**
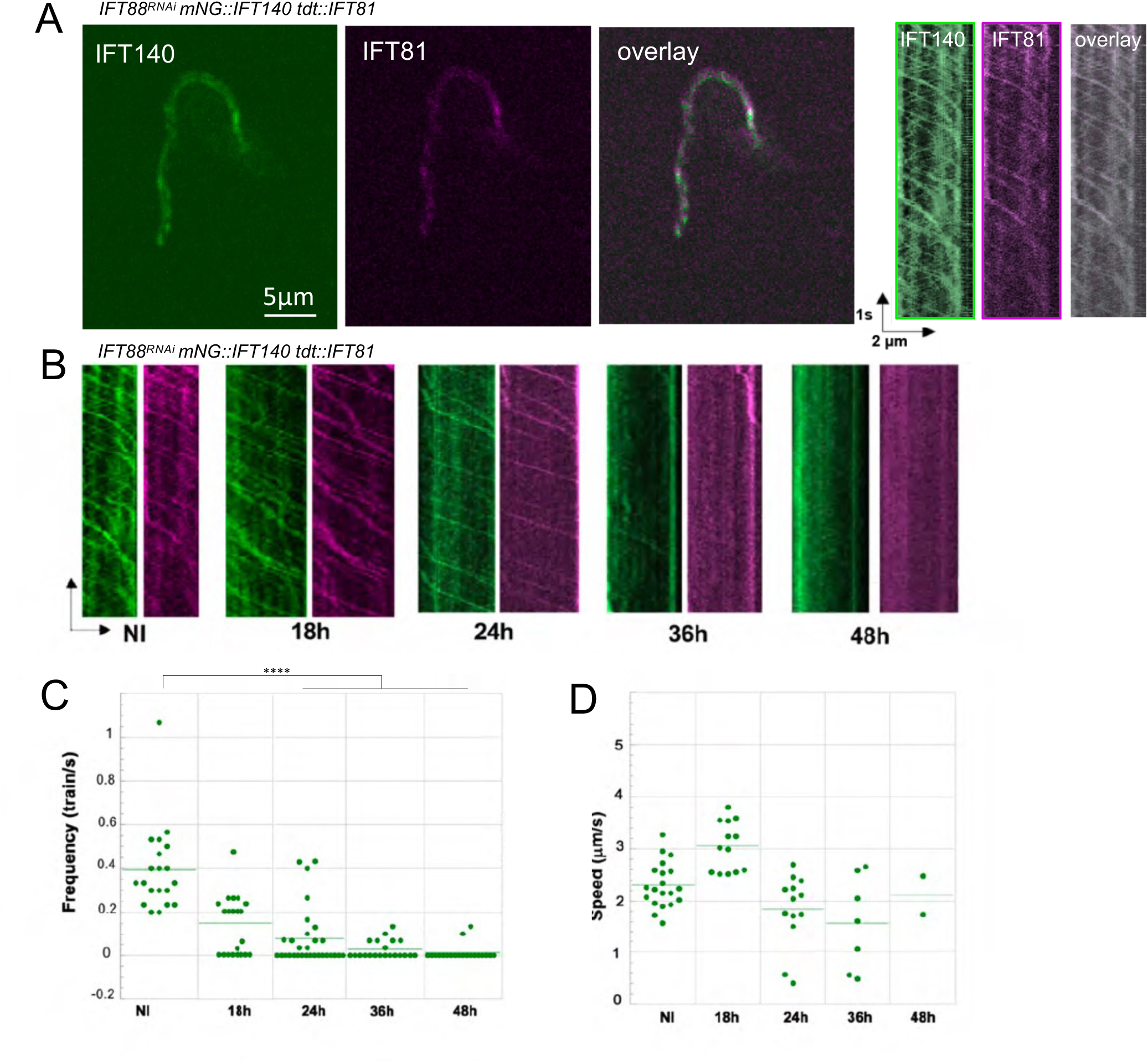
IFT88 is required for intraflagellar transport of IFT-A and IFT-B reporters. (A) Still images of an *IFT88^RNAi^* cell expressing mNG::IFT140 (green) and tdT::IFT81 (magenta) observed with a spinning disk microscope equipped with two cameras. The overlay shows the merged mNG and tdT images. Corresponding kymographs are showed on the right panels with the same colour code. (B) Kymographs obtained from *the IFT88^RNAi^ mNG::IFT140 TdT::IFT81* cell line during the course of RNAi induction (NI, RNAi non-induced). Kymographs show mNG::IFT140 (green) and tdT::IFT81 (pink). (C) Graph representing the frequency of anterograde trains observed with the *IFT88^RNAi^ mNG::IFT140 TdT::IFT81* cell line during the course of induction. Each point represents the average of 20 cells. (D) Graph representing the speed of anterograde trains observed for the same trypanosomes of the *IFT88^RNAi^ mNG::IFT140 TdT::IFT81* cell line. Each point represents the average frequency and speed of all detected trains in 20 different cells. ****p<0.0001(Anova test)

**Figure 5.**
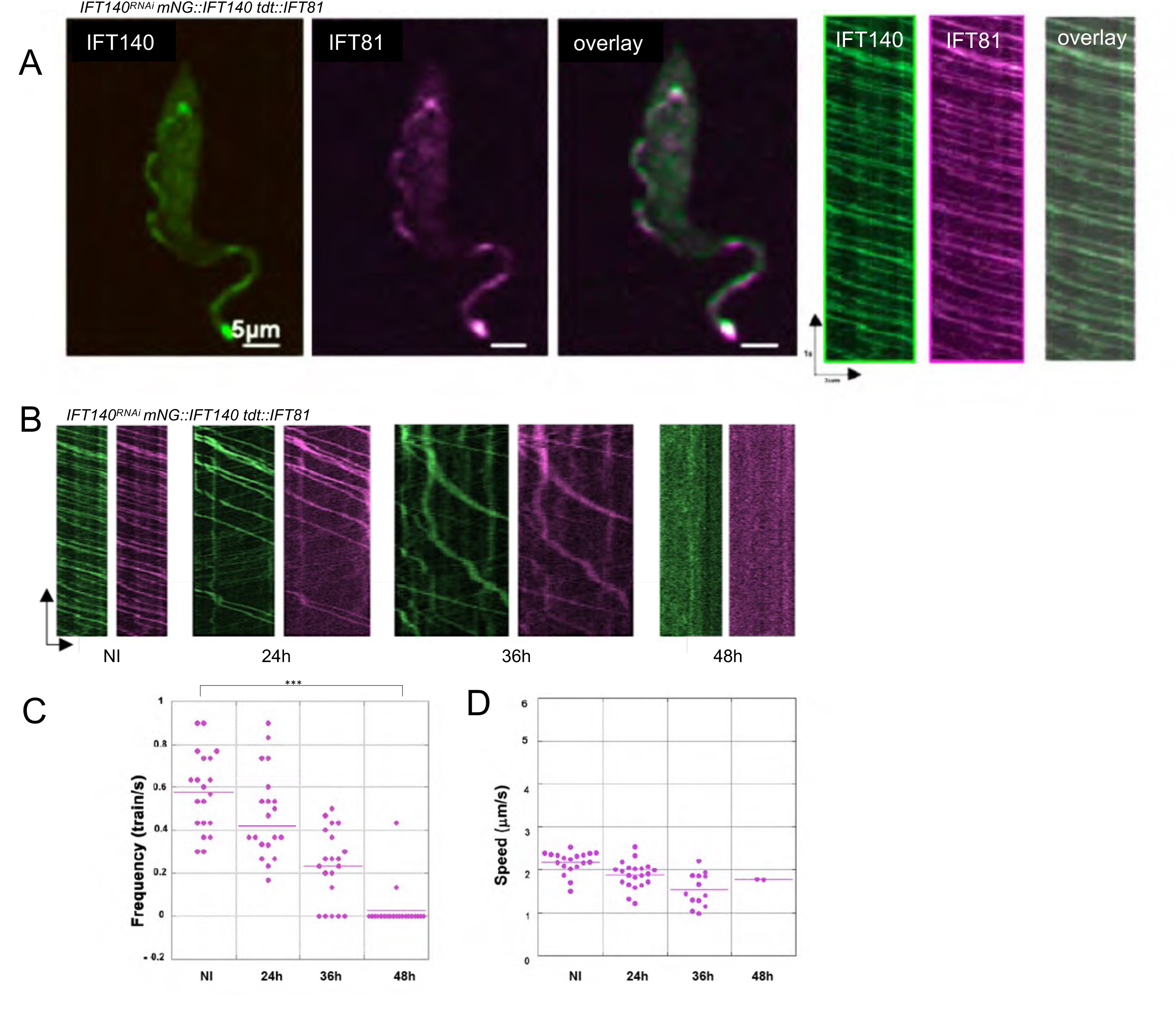
IFT140 is essential for trafficking of IFT88. (A) Still images of *IFT140^RNAi^*cells expressing mNG::IFT140 (green) and tdT::IFT81 (magenta) observed with a spinning disk microscope equipped with two cameras. The overlay shows the merged mNG and tdT images. Corresponding kymographs are shown on the right panels with the same colour code. (B) Representative kymographs obtained from the *IFT140^RNAi^* cell line expressing mNG::IFT140 (green) and TdT::IFT81 (pink) at the indicated times of induction (NI, RNAi non-induced). IFT frequency is reduced but remaining trains still contain both mNG::IFT140 and tdT::IFT81. (C) Frequency and (D) speed of anterograde IFT trains are represented on graphs. Each point represents the average frequency and speed of all detected trains in 20 different cells. *** p<0.001 (Anova test)

### IFT88 is essential for assembly of anterograde IFT trains

The IFT-B complex is defined as the central element of IFT trains (Jordan et al., 2018). To formally evaluate its role in the construction of IFT trains in trypanosomes, we studied the *IFT88^RNAi^ mNG::IFT140 TdT::IFT81* strain. Four knockdown induction times (18h, 24h, 36h and 48h) were collected in addition to the non-induced control (Fig. 4B). In control conditions, fluorescent signals for mNG::IFT140 and tdT::IFT81 colocalize, as described above (Fig. 4B, NI). After induction of RNAi by addition of tetracycline, we observed the expected cellular phenotypes: inhibition of new flagellum formation, emergence of small non-flagellated cells while old flagella assembled before RNAi remained present and their length was maintained, as previously reported (Fort et al., 2016; Kohl et al., 2003). As early as 18 or 24h after initiation of RNAi, the movement of both mNG::IFT140 and tdT::IFT81 was affected, with a drop in IFT train frequency and a reduction of the intensity of the fluorescent traces for the remaining trains, affecting equally both proteins that nevertheless remain colocalised at all induction times (Fig. 4B). After 48h in RNAi conditions, most cells did not exhibit IFT trafficking and those that did displayed very low frequency (only one train every 10 seconds, Fig. 4B-C). Train speed was less affected, with yet a curious increase at the 18h time-point and a slight decrease later on (Fig. 4D). These results demonstrate that anterograde IFT trafficking is severely impacted by the depletion of IFT88 and that IFT-A proteins cannot travel independently, in agreement with the proposed central role of the IFT-B complex for train formation.

To evaluate the impact of IFT88 knockdown on the length of IFT trains in flagella, we turned towards volumetric electron microscopy in both uninduced conditions and at 18h and 48h in RNAi conditions. Samples were fixed before being dehydrated and embedded in resin, then imaged in FIB-SEM. The volumes obtained have an isotopic resolution of 10nm, giving the possibility to monitor IFT trains all along flagella as these can appear in any orientation within the volume. Electron-dense particles located between the flagellar membrane and microtubule doublets were detected, segmented and measured with MatLab and R software as described previously (Bertiaux et al., 2018a). Original volumes containing segmented flagella and IFT trains are presented at Video S4 (left panels). In non-induced *IFT88^RNAi^* cells (Fig. 6A), a total of 54 trains were detected in 6 different portions of flagella (Table 2). All these trains were located close to microtubule doublets 3-4 and 7-8, as in wild-type cells (Bertiaux et al., 2018a). Trains displayed a mixture of lengths, ranging from less than 100 nm to close to 3 µm (Fig. 7A, grey bars). Observation of *IFT88^RNAi^* samples induced for 18h showed a pronounced reduction in train length compared to non-induced samples (Fig. 6B, Video S4), not exceeding 500 nm with one exception (Fig. 7A, orange bars). This drop was more severe after 48h of IFT88 knockdown where all the remaining trains were only 200 nm or shorter (Fig. 7A, blue bars). Overall, the trains occupied barely 4% of the total axoneme length, in contrast to 32% in non-induced conditions (Table 2). Therefore, progressive depletion of IFT-B results in a spectacular decrease of the length of IFT trains. We conclude that the IFT-B complex is rate-limiting for the assembly of anterograde IFT trains.

**Figure 6.**
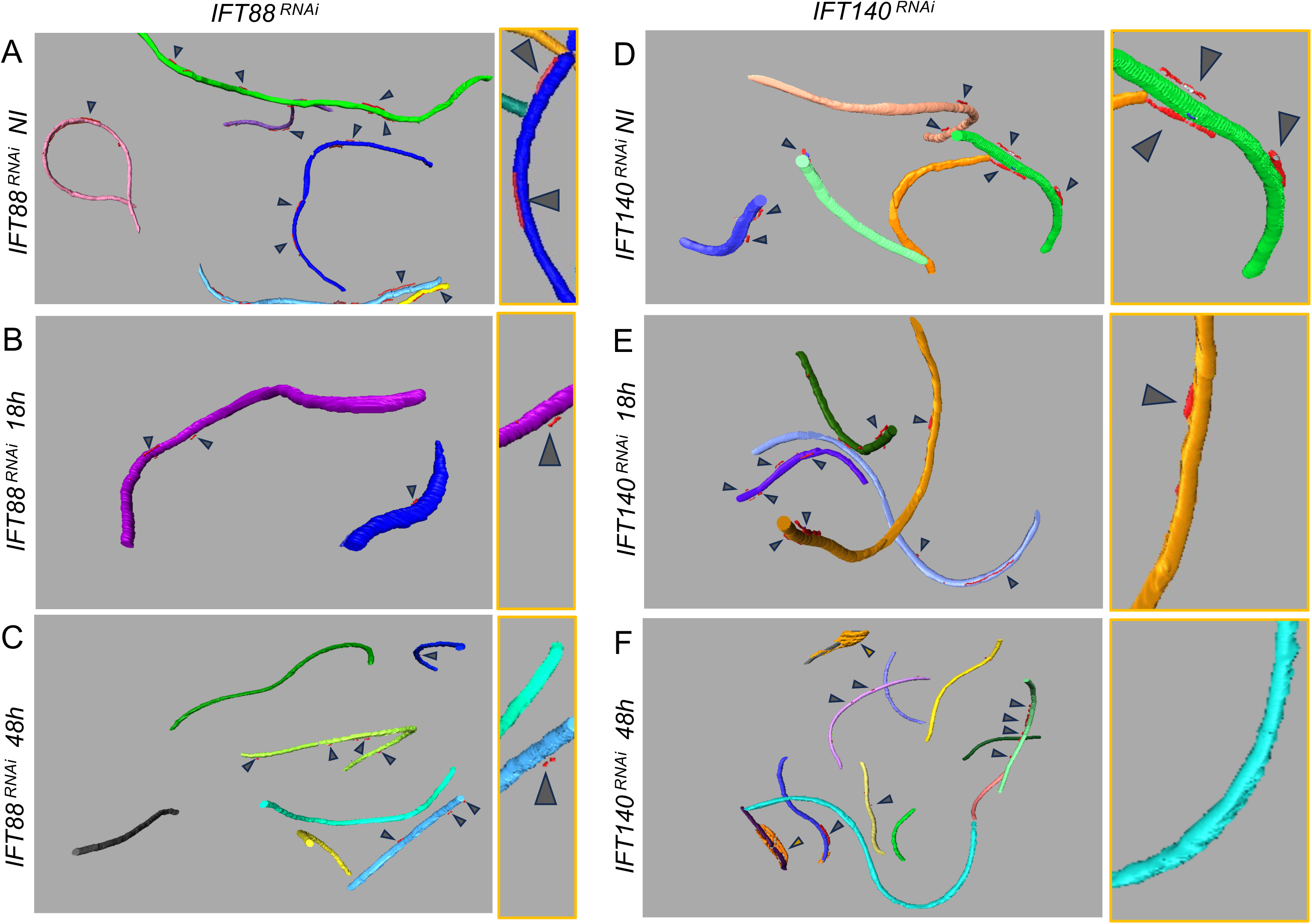
IFT88 and IFT140 are required for IFT train construction. FIB-SEM data obtained after segmentation of different flagella (each represented in a different colour) showing the presence of multiple IFT trains (in red, indicated with dark grey arrows) from the *IFT88^RNAi^* (A-C) or the *IFT140^RNAi^* (D-F) cell line in non-induced conditions (A,D), after 18h (B,E) and 48h (C,F) in RNAi conditions. A zoom on a flagellum is displayed on the right-side of each panel. Original FIB-SEM images and segmented flagella are presented at Video S4 (left panels) and detailed quantification is shown at Table 2.

**Figure 7.**
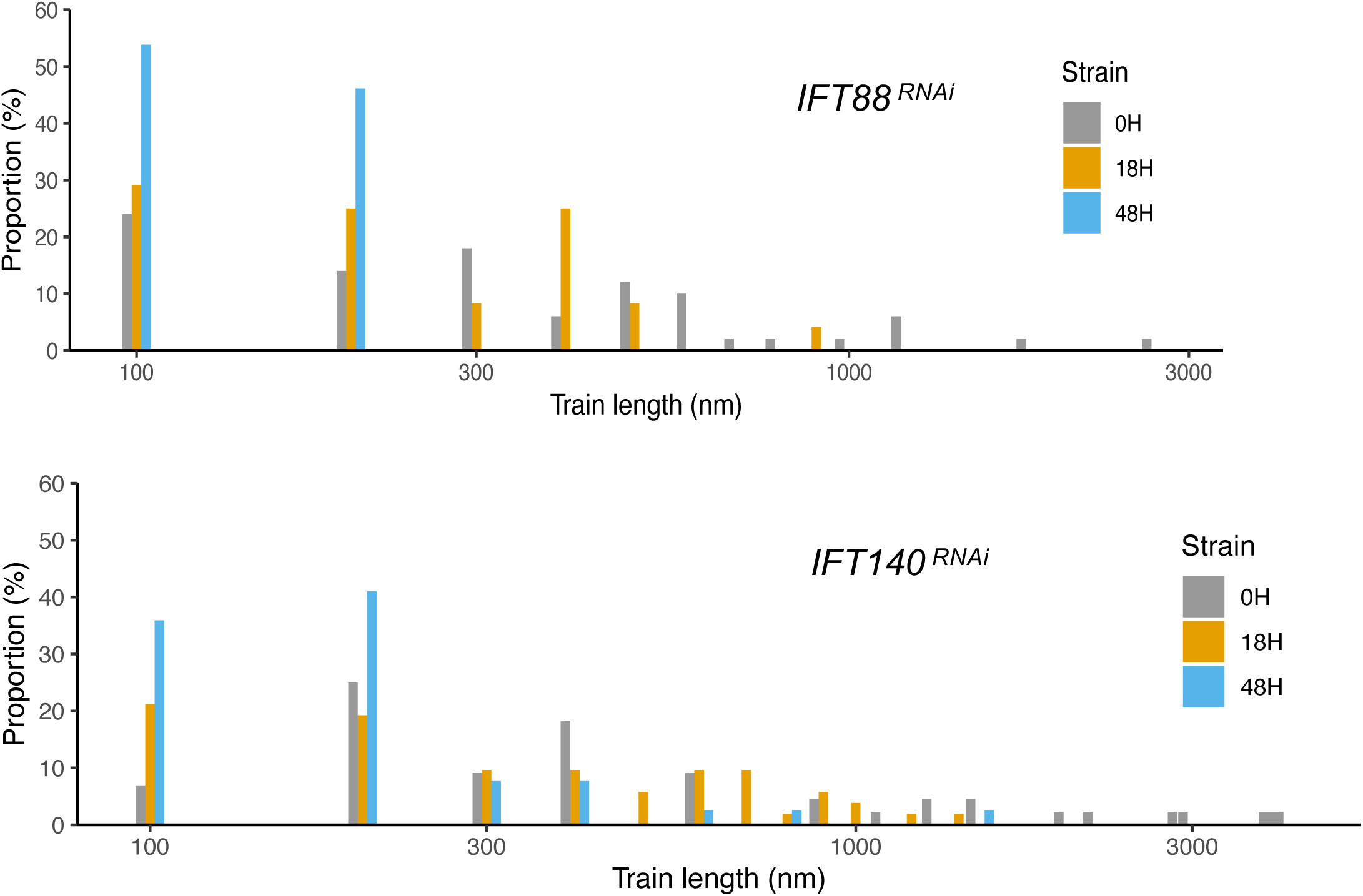
The length of IFT trains depends on both IFT-A and IFT-B presence. Length distribution (log scale) of IFT trains observed for *IFT88^RNAi^* (A) and *IFT140^RNAi^* (B) by FIB-SEM in non-induced conditions (grey bars), after 18h (orange bars) and 48h of induction (blue bars). A clear shift in train length is visible during the course of both knockdown.

**Table 2.** IFT train lengths measured in FIB-SEM samples of *IFT88^RNAi^* and*IFT140^RNAi^* cell lines after different times of induction. Nb of flagella: number of flagella analysed; Cumulated length of axoneme: total length of all axonemes analysed in a volume; Nb of IFT trains: number of IFT trains detected on these flagella; IFT train length: average length of all measured IFT trains; Frequency: number of IFT trains detected per µm (independent of their length); Cumulated length of IFT trains: the sum of the lengths of all IFT trains analysed; Axoneme occupancy: percentage of axoneme occupied by IFT trains.

| *IFT88<sup>RNAi</sup>* | Nb of flagella | Nb of IFT trains | Frequency (nb train/μm) | Cumulated length of axoneme (μm) | Cumulated length of IFT trains (μm) | IFT trains length (nm) | % Axoneme occupancy by IFT trains |
| --- | --- | --- | --- | --- | --- | --- | --- |
| NI | 6 | 54 | 0.75 | 72 | 22.9 | 420 | 32 |
| 18h | 11 | 32 | 0.82 | 39 | 5.7 | 196 | 14.6 |
| 48h | 7 | 13 | 0.39 | 33 | 1.4 | 115 | 4 |

| *IFT140<sup>RNAi</sup>* | Nb of flagella | Nb of IFT trains | Frequency (nb train/μm) | Cumulated length of axoneme (μm) | Cumulated length of IFT trains (μm) | Length of IFT trains (nm) | % Axoneme occupancy by IFT trains |
| --- | --- | --- | --- | --- | --- | --- | --- |
| NI | 16 | 47 | 0.51 | 90 | 38 | 1087 | 42 |
| 18h | 11 | 59 | 0.66 | 71 | 20 | 341 | 28 |
| 48h | 16 | 41 | 0.32 | 151 | 8 | 398 | 5 |

### IFT140 is also essential for the formation of anterograde trains

In a few cases, IFT-A proteins can form trains independently of the presence of IFT-B (Ou et al., 2005; Toriyama et al., 2016). To assess the contribution of IFT-A in formation of anterograde trains in trypanosomes, we used the cell line expressing simultaneously the same mNG::IFT140 and TdT::IFT81 fusion proteins but this time in the *IFT140^RNAi^* strain. In non-induced controls, both proteins displayed robust IFT and colocalise in live cells (Video S3), as said above (Fig. 5A). The impact of inhibiting IFT140 abundance on the transport of TdT::IFT81 was monitored over time upon RNAi knockdown from 0 to 48 h (Fig. 5B). As expected, the number of moving trains tagged with mNG::IFT140 decreased with induction time, which was clearly visible on the kymographs (Fig. 5B, green channel), confirming RNAi efficiency. Remarkably, the same reduction in the frequency of tdT::IFT81 transport events was observed at all induction times (Fig. 5B, magenta channel). The latter was not detected traveling alone, ruling out the possibility that IFT-B proteins could form moving trains in the absence of IFT-A. In fact, there was still >90% co-localisation of the mNG::IFT140 and tdT::IFT81 signals at induction times of 24h and 36h (see kymographs, Fig. 5B). After 48 h of induction, there was no longer TdT::IFT81 movement, with a couple of exceptions where frequency was much reduced (Fig. 5B-C). A mild decrease in IFT anterograde speed was observed during the course of induction (Fig. 5D).

Three *IFT140^RNAi^* samples were prepared for FIB-SEM (uninduced condition, after 18h and 48h of induction, Video S4, right panels). Flagella and IFT trains were segmented and measured as above. In non-induced conditions, a similar profile to non-induced *IFT88^RNAi^*cells was observed (Fig. 7A, grey bars), with a mixture of short and long trains, some of them reaching up to 4 µm (Fig. 7B, grey bars). After 18 hours of RNAi induction targeting IFT140, the length of IFT trains was reduced by more than half, with only a couple of trains a bit longer than 1 µm (Fig. 7D orange bars and Table 2). After 48 hours, much fewer trains were detected, and their length appeared even smaller (Fig. 7B, blue bars and Table 2). This was confirmed when the cumulative percentage occupancy of IFT trains over total axoneme length was calculated. In the uninduced strain, IFT trains occupied 42% of total axoneme length, in line with their frequency of detection in conventional electron microscopy sections (Fort et al., 2016). On the contrary, they occupied 28% after 18h of induction and only 5% after 48h of induction (Table 2). This drastic drop can be explained by the decrease in train length and frequency and is comparable to that observed in the *IFT88^RNAi^* strain (Table 2) where IFT-B was targeted.

We conclude that the IFT-A complex is essential for anterograde train construction, since reducing IFT140 abundance affects train quantity and length in a very similar way to what was observed for IFT88 knockdown, albeit with a slower kinetics.

## Discussion

In the literature, two different phenotypes have been reported upon perturbation of IFT-A expression. When IFT-A is absent or greatly reduced, flagellum assembly is blocked, with little or no axonemal microtubules, as observed in null mutants in *Chlamydomonas* (Picariello et al., 2019; Zhu et al., 2017) and *Leishmania* (Sunter et al., 2018). However, when low amounts of IFT-A remain present, either in RNAi experiments or in the case of mutant proteins, shorter cilia or flagella are assembled and their extremities are filled with IFT proteins. This has been reported for *Chlamydomonas* (Iomini et al., 2009; Jordan et al., 2018) and *C. elegans* (Blacque et al., 2006; Efimenko et al., 2006) as well as in human patients or mammalian cell lines (Arts et al., 2011; Bredrup et al., 2011; Davis et al., 2011; Mill et al., 2011; Perrault et al., 2012).

Both situations were encountered in *T. brucei* following RNAi against IFT140 or IFT122, but in a time-dependent manner: at early stages of knockdown, cells assemble short flagella with accumulation of IFT-B proteins at their tip while at advanced stages, new flagella are not constructed anymore (Absalon et al., 2008a; Absalon et al., 2008b; Blisnick et al., 2014). This prompted us to revisit the role of the IFT-A complex in anterograde train formation, taking advantage of the stability of trypanosome flagella assembled before the onset of RNAi. This allowed us to visualise anterograde trains in existing flagella both in live and fixed cells, and to measure their length using FIB-SEM. In normal conditions, IFT-A, IFT-B and dyneins exhibit the expected ratio (here based on knowledge acquired in cryo-EM studies of trains rin *Chlamydomonas*)(Fig. 8A). At early time points of RNAi knockdown, when residual amounts of IFT140 were still present, moving trains were detected but their frequency and their length were reduced (Fig. 8B-C). At advanced stages of knockdown, when little IFT140 protein remained present, trains were hardly detected (Fig. 8D). We conclude that the IFT-A complex is required for the efficient formation of anterograde trains and its absence explains the no-flagellum phenotype reported in several circumstances since flagella cannot be constructed without IFT (Picariello et al., 2019; Sunter et al., 2018; Zhu et al., 2017). This essential contribution of the IFT-A complex to efficient train formation could be due to its positioning in the edifice where it appears to bridge two IFT-B complexes together and so could facilitate IFT-B multimerization or stabilisation (Jordan et al., 2018; Lacey et al., 2023; van den Hoek et al., 2022).

**Figure 8.**
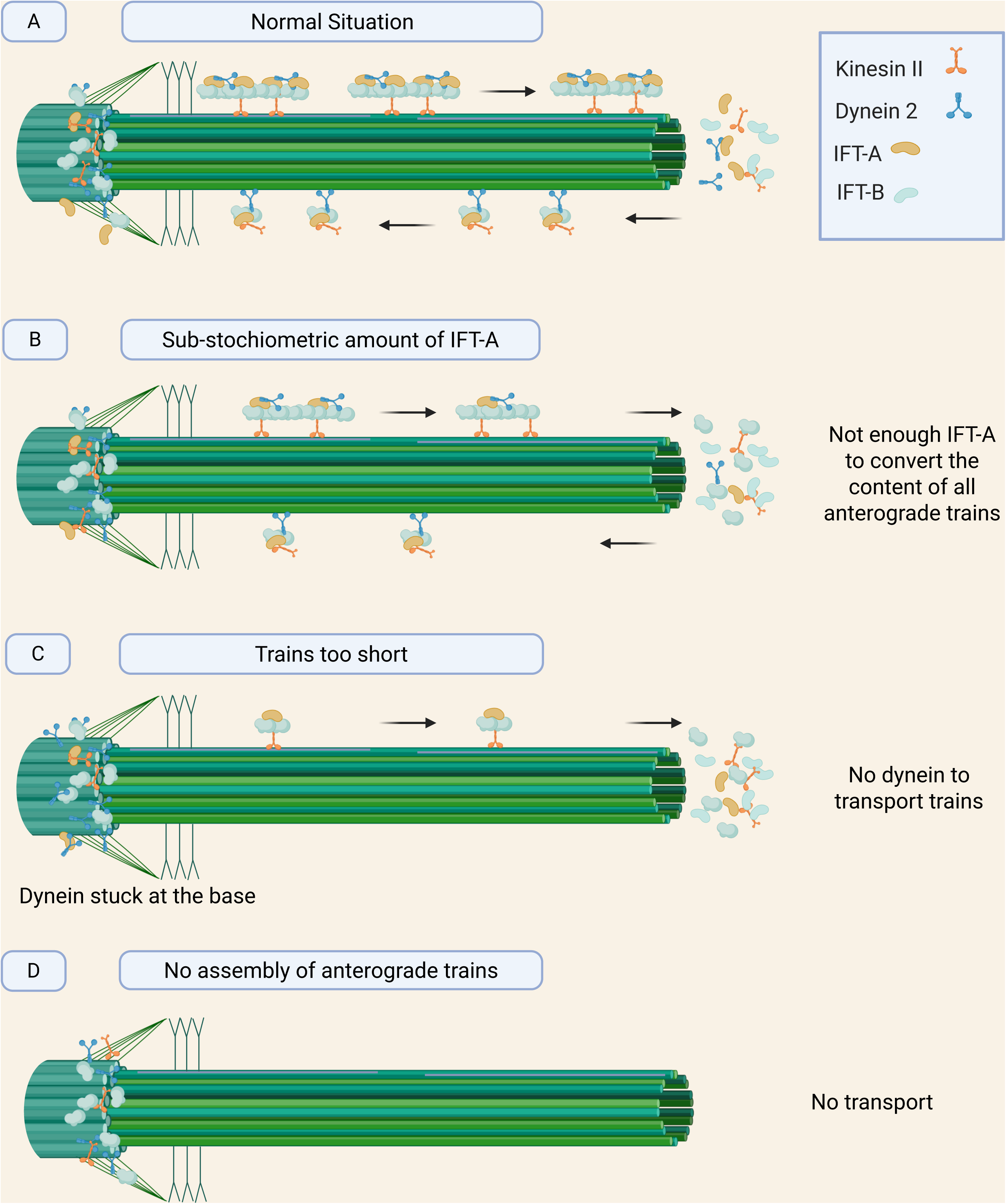
IFT-A is required for proper assembly of anterograde IFT trains. (A) In the normal situation, IFT proteins and motors concentrate at the base of the flagellum at the level of transition fibres (Araujo Alves et al., 2025). Anterograde trains composed of IFT-B complexes (green), IFT-A complexes (yellow) and dynein (blue) in a predicted 8:4:2 stoechiometry (Jordan et al., 2018) circulate towards the tip driven by the kinesin motor (orange). The actual amounts of kinesin motors per train are unknown, but mass spectrometry analysis of purified flagella suggests a lower concentration compared to other components (Subota et al., 2014). At the tip, trains are split in shorter retrograde trains driven by the dynein motor. In *T. brucei*, the kinesin motor is recycled by retrograde transport (Alves et al., BioRXiv 2026), as also observed in *C. elegans* (Prevo et al., 2015). (B-C) When the amount of IFT-A is reduced, shorter anterograde trains are assembled and this can have two consequences: either the proportion of IFT-A in these trains is too low relative to IFT-B complexes, which would mean an unsufficient number of IFT-A at the tip to ensure conversion of all the material in retrograde trains, hence resulting in accumulation of IFT-B proteins (B). Alternatively, trains might have the right proportion of IFT-A and IFT-B but their short size would not be sufficient to load dynein that would remain at the base of the organelle (C). In absence of dynein, even if anterograde trains were able to convert to retrograde trains, they would not be transported without the motor, also resulting in accumulation of material at the tip. (D) Finally, if IFT-A proteins are absent, IFT-B alone cannot make efficient anterograde trains and IFT is arrested.

Our discovery that trains become shorter upon reduction of the IFT-A amount resonates with at least two previous studies and could provide an explanation to the frequently reported retrograde phenotypes. First, in the seminal cryo-electron tomography paper defining anterograde IFT train anatomy, Jordan et al. (2018) used the *Chlamydomonas ift139-2* cell line. The mutation in IFT139 led to a 10-fold drop in the flagellar amount of IFT-A proteins but yet anterograde trains were still assembled, although they were shorter than in control cells (193 nm vs 310 nm) and appeared distorted. Densities corresponding to IFT-A complexes were detected but they were found to be sub-stoichiometric relative to IFT-B and often not visible after sub-tomogram averaging. A few trains made of only IFT-B were also mentioned. This *ift139-2* cell line assembles flagella but these display accumulation of material at their distal tip (Jordan et al., 2018), suggesting defects in IFT recycling. In another study, a *Chlamydomonas* cell line expressing a mutant version of IFT140 lacking all its WD repeats exhibited reduction in anterograde IFT trafficking, but 2-colour live imaging revealed that all the trains were containing IFT140 and IFT20 (a reporter for the IFT-B complex), suggesting that the presence of at least low doses of IFT-A is required for anterograde trains. Similarly, the vast majority of trains observed in our trypanosome study contained both IFT140 (IFT-A) and IFT81 (IFT-B).

We propose that in conditions where the level of IFT-A protein is reduced (but not missing), shorter anterograde trains are assembled with potentially two types of defects. First, trains could contain lower than normal amounts of IFT-A complex relative to IFT-B (Jordan et al., 2018). Since conversion of anterograde train to retrograde train depends on the IFT-A complex (Goncalves-Santos et al., 2023; Lacey et al., 2024), a sub-stoichiometric ratio of IFT-A would mean that not all the IFT-B complexes can be recycled (Fig. 8B), even if dynein still associates normally to the IFT-B complex (Jordan et al., 2018). Second, anterograde trains with low amounts of IFT-A might be too short to efficiently load dynein since the expected ratio is 8 copies of the IFT-B complex for 4 copies of the IFT-A and 2 copies of dynein (Jordan et al., 2018). For example, a train composed of 2 copies of IFT-B and 1 copy of IFT-A is unlikely to load dynein. So, even if it converted to retrograde trains, transport would be impossible due to the absence of motor (Fig. 8C). If dynein is not efficiently associated to IFT trains, it should be missing from or found in reduced amounts in short bulbous flagella when IFT-A proteins are depleted. This was indeed the case in the trypanosome IFT140 RNAi knockdown in *T. brucei*: the heavy chain DHC1b and the light intermediate chain DLI1 (also known as XBX-1) remained at the base (Blisnick et al., 2014). The same phenotype was reported for XBX-1 in the *IFT140* mutant of *C. elegans* (called *che-11*)(Schafer et al., 2003).

These results were logically interpreted as a retrograde transport phenotype but in fact the retrograde defect would be a secondary consequence of the construction of aberrant anterograde trains (Fig. 8B-C). This explanation is compatible with mutant proteins reported in multiple human patients that function less well but could still sustain formation of abnormally short IFT trains. Modelling of these mutations has shown that those present at the TPR regions would affect train formation or stability, although possible impact on cargo and membrane interaction can also be considered (Lacey et al., 2023). In the mutant *Chlamydomonas* IFT140 protein mentioned above, the frequency of retrograde transport was more affected than that of anterograde transport (Picariello et al., 2019), in agreement with defective train recycling.

In conclusion, the unique features of the trypanosome model allowed the clarification of the contribution of the IFT-A complex in anterograde train assembly, in addition to its well-recognised role in retrograde train formation. Combined to structural data reported in *Chlamydomonas* (Jordan et al., 2018; Lacey et al., 2023; Lacey et al., 2024), this finding resolves the conundrum of why inhibition of IFT-A proteins sometimes results in the absence of cilia or in other occasions in the construction of short cilia filled with IFT particles.

## Material & Methods

### Trypanosome cell lines and cultures

Cell lines used for this work were derivatives of *T. brucei* strain 427 cultured at 27°C and in SDM79 medium supplemented with hemin and 10% foetal calf serum. The 29-13 cell line expresses the T7 RNA polymerase and the tetracycline repressor, allowing for tetracycline-inducible expression (Wirtz et al., 1999). It was transformed to express complementary single-stranded RNA corresponding to a fragment of the *IFT140* or *IFT88* gene from inducible T7 promoters facing each other in the pZJM vector following integration in the rDNA locus of cells (Absalon et al., 2008b; Kohl et al., 2003; Wang et al., 2000). Addition of tetracycline (1µg/ml) to the medium induces expression of sense and anti-sense RNA strands that can anneal to form double stranded RNA and trigger RNAi. Two approaches were used for *in situ* tagging of IFT proteins with fluorescent reporters. For IFT81 and IFT140, the first 500 nucleotides of the corresponding genes were chemically synthetized (GeneCust, Luxembourg) and cloned in frame with the mNeonGreen or the TandemTomato gene within the p2675 vector (Kelly et al., 2007). Plasmids were linearised within the gene of interest and introduced by nucleofection in the described cell lines. For IFT122 and IFT139, the long primer PCR tagging strategy was used with the pPOTv7 vector (Dean et al., 2015). Clonal populations were obtained by limiting dilution. Cell lines were characterized either by phase contrast or direct fluorescence observation (mNeonGreen or tandemTomato). Cell culture growth was monitored daily with an automatic Muse cell analyser (Merk Millipore, Paris).

### Single and dual colours live cell imaging of IFT trafficking

Cell lines expressing mNG or tdT fusion proteins were grown in standard conditions and samples were mounted between glass and coverslip for observation on a Perkin Elmer UltraView Vox microscope equipped with a Yokagawa CSUX1 spinning- disk, and two cameras for dual-colour simultaneous imaging (Bertiaux et al., 2018a). mNG was excited with a 488 nm laser (500 mW, SAPPHIRE 488-500, Coherent) and detected through an emission filter (BP 500-550 nm) while tdT was excited with a 554 nm laser and detected through an emission filter (581nm). For dual colour imaging, a dichroic mirror splits the signal on two Hamamatsu EM-CCD cameras operating simultaneously and videos were acquired with an objective 63x with 1.4NA and the Volocity software. Cells were maintained at 27°C under a temperature-controlled chamber. Sequences of 30 seconds were acquired with an exposure time of 100 ms per image. Kymographs of IFT anterograde trains were extracted using the Kymograph tracker 2 with the Icy software (de Chaumont et al., 2012). Graph representations were done with the Kaleidagraph software V4-5-2. Statistical analyses were done with Prism v10.0 using Anova test. This test was selected to detect if there were statistically significant differences between the means, knowing that variance of each mean appeared similar. Statistically significant differences are indicated with two stars (p<0.05), three stars (p<0.001 or four stars (p<0.0001).

### Focused Ion Beam - Scanning Electron Microscopy (FIB-SEM)

Trypanosomes were fixed directly in medium with a final concentration of 2.5% glutaraldehyde (Sigma), cells were spun down, the supernatant was discarded, and the pellet was fixed and treated exactly as before (Bertiaux et al., 2018a). IFT trains were defined as electron dense structures sandwiched between the axoneme and the flagellum membrane and present on a minimum of 3 consecutive slices (30 nm). Densities associated to membrane distortions were excluded. Trains were defined as different when separated by a minimum of 3 slices (30 nm).

### Data processing, 3-D-reconstruction, and length measurement of IFT trains

Alignment of image stacks was done with the open-source software Fiji for data alignment (Schindelin et al., 2012) and Amira Software for visualization (FEI Thermofisher, v6.0.1). Segmentation and 3-D reconstructions were performed semi-automatically using Amira software and were corrected manually. Segmentations of flagella and IFT trains were first split according to their segmented colours then skeletonized in Fiji (Schindelin et al., 2012) using the Skeletonize 3D plugin (Arganda-Carreras et al., 2010). The number of voxels composing the generated skeletons was computed in ImageJ using the Object Counter 3D plugin (Bolte and Cordelieres, 2006). The length of analysed biological structures (flagella and IFT trains) was calculated as the number of voxels constituting the skeletonized structures multiplied by the size of the voxel. IFT train data (Fig. 7) was generated with the software R (R Core Team, 2014).

## Acknowledgements

We are grateful for Ultrastructural BioImaging unit support by the GIS-IBISA, the French Government Programme Investissements d’Avenir France BioImaging (ANR-24–INBS–0005 FBI BIOGEN), and the DIM1Health program. We also thank the Photonic BioImaging unit for their advice and access to the UltraVIEW VOX system. Special thanks to Jean-Yves Tinevez for his guidance and training on the spinning disk microscope. We thank Cyril Scandola from UBI for help with statistical analyses with Prism software. E.B. and C.F. were supported by fellowships from the French National Ministry for Research and Technology (doctoral school CDV515) and from La Fondation pour la Recherche Médicale (FDT20170436836 and FDT20150532023). Research is funded by grants from the ANR (ANR-18-CE13-0014-01), the Fondation pour la Recherche Médicale (FRM-EQU202203014654) and the French Government Investissement d’Avenir programme, Laboratoire d’Excellence “Integrative Biology of Emerging Infectious Diseases” (ANR-10-LABX-62-IBEID).

The authors declare that they have no conflict of interest.

## Author’s contributions

A.M.: Conceptualisation, data curation, formal analysis, investigation, visualisation, writing-original draft, writing-review and editing; T.B.: Conceptualisation, formal analysis, investigation, supervision, visualisation, writing-review and editing; E.B.: Investigation, writing-review and editing; C.F., investigation, writing-review and editing; M.M.: data curation, visualisation; S.T.: data curation, formal analysis, methodology, visualisation, writing-review and editing; P.B.: Conceptualisation, funding acquisition, project administration, supervision, writing-original draft, writing-review and editing.

The authors declare no competing financial interests.

